# Regulatory RNA design through evolutionary computation and strand displacement

**DOI:** 10.1101/004580

**Authors:** William Rostain, Thomas E. Landrain, Guillermo Rodrigo, Alfonso Jaramillo

## Abstract

The discovery and study of a vast number of regulatory RNAs in all kingdoms of life over the past decades has allowed the design of new synthetic RNAs that can regulate gene expression *in vivo.* Riboregulators, in particular, have been used to activate or repress gene expression. However, to accelerate and scale up the design process, synthetic biologists require computer-assisted design tools, without which riboregulator engineering will remain a case-by-case design process requiring expert attention. Recently, the design of RNA circuits by evolutionary computation and adapting strand displacement techniques from nanotechnology has proven to be suited to the automated generation of DNA sequences implementing regulatory RNA systems in bacteria. Herein, we present our method to carry out such evolutionary design and how to use it to create various types of riboregulators, allowing the systematic *de novo* design of genetic control systems in synthetic biology.

## 1 Introduction

RNA is a versatile molecule, and its many roles in the cell include enzyme-like activity and regulation, in addition to its various roles in translation [1]. Some of RNA’s many mechanisms for modulating biological processes have been harnessed by synthetic biologists for creating artificial regulators of gene expression [2, 3]. Examples in bacteria or yeast involve positive and negative riboregulators [4–9], ribozyme-based systems [10–13], or CRISPR systems [14–16]. These regulatory RNAs can control gene expression through base pairing with a messenger RNA (mRNA) or DNA, and typically have a defined secondary structure that ensures stability and functionality (mainly for interaction).

In this work, we focus on bacterial small RNAs (sRNAs) that can interact with the 5’ untranslated region (5’ UTR) of a given mRNA (see Table 1). Using computational tools, from which RNA secondary structures and base pairing energies and probabilities can be predicted [17, 18], in combination with *in silico* evolutionary design principles [19, 20], various RNA systems can be engineered. Although synthetic RNAs could be designed *by hand*, fast and large-scale engineering of complex RNA circuits for synthetic biology cannot be based on this lengthy, case-by-case design strategy. To attain this goal, computer-assisted design will be required to accelerate the design process [7, 12].

**Table 1:**
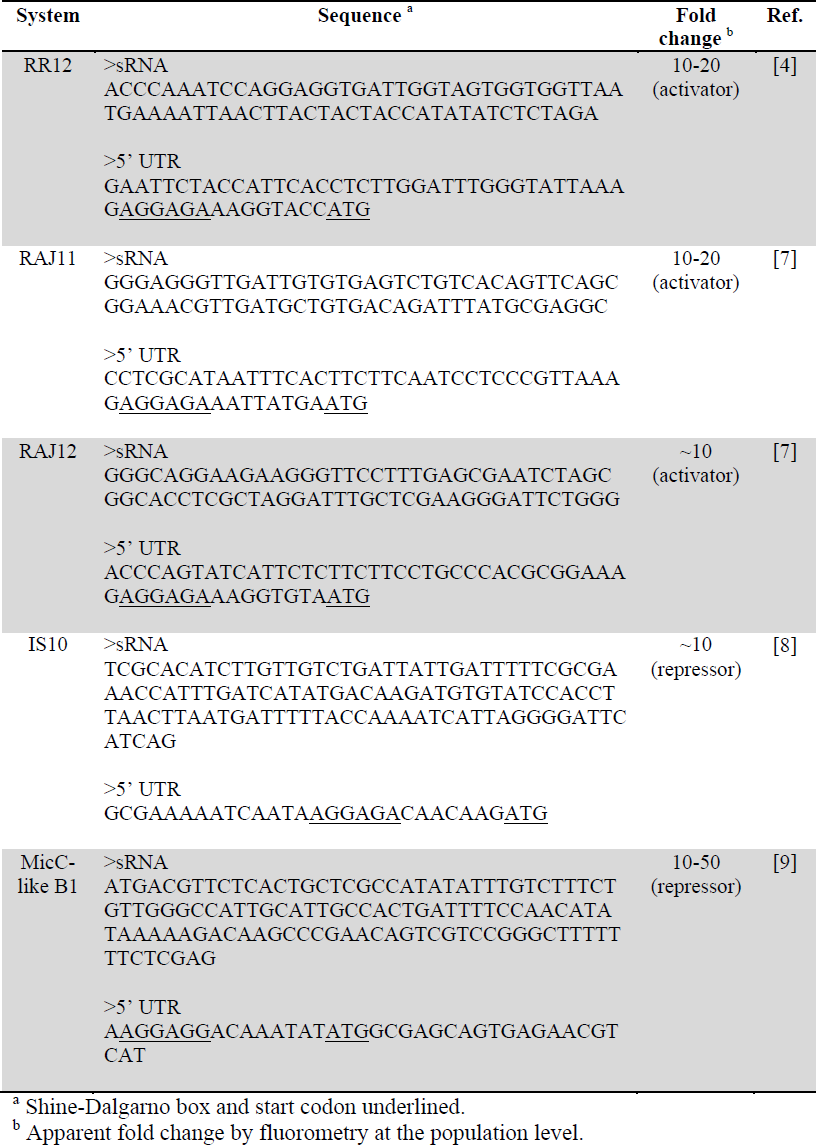
List of engineered riboregulators of gene expression.

Automated design has been successful for some types of RNAs, as well as for nucleic acids in general, proteins and circuits (applications reviewed in ref. [19]), allowing the diffusion of these methods as general tools for molecular biology. Evolutionary computation of riboregulation starts with two RNA sequences and iterates alternative rounds of mutation and selection of species that show the desired structural characteristics. This method adapts strand displacement techniques from nanotechnology and it has successfully been applied to the *de novo* design of bacterial riboregulators [7]. It has the advantage of being based on physiochemical principles, assessing all specifications and constraints of the design processs *in silico* and without manual inspection, and it can be tailored to the design of simple or complex riboregulation [21].

In the following pages, we detail the strategy used for creating a software capable of performing the evolutionary computation of two interacting sRNAs. This involves creating an objective function used to score RNA sequences and to determine whether they will show the desired behavior, together with a mutation operator used to efficiently search the sequence space. We then explain how to use this method to create positive and negative riboregulation [2], providing along the way some tips and resources for the design, implementation and characterization of such systems. We expect this approach will be of value for engineering genetic circuits *in vivo* and for increasing our ability to design more sophisticated regulatory systems.

## 2 Materials

### 2.1 Software

To computationally design regulatory RNA systems, which perform a particular logical operation, a combinatorial optimization algorithm is constructed, in which thermodynamic and structural parameters are used to evaluate a system at any time during the optimization (see Fig. 1). To estimate those parameters, the ViennaRNA package [22] is used. The different intra-and intermolecular secondary structures and the corresponding free energies are thus easily calculated throughout the evolutionary process (see section 3.1.2). Once the solutions have converged, the Nupack package [23] is used to carry out a post-analysis of the designed sequences. This independent software serves as a subsequent filtering and validation tool (see section 3.1.4).

**Figure 1:**
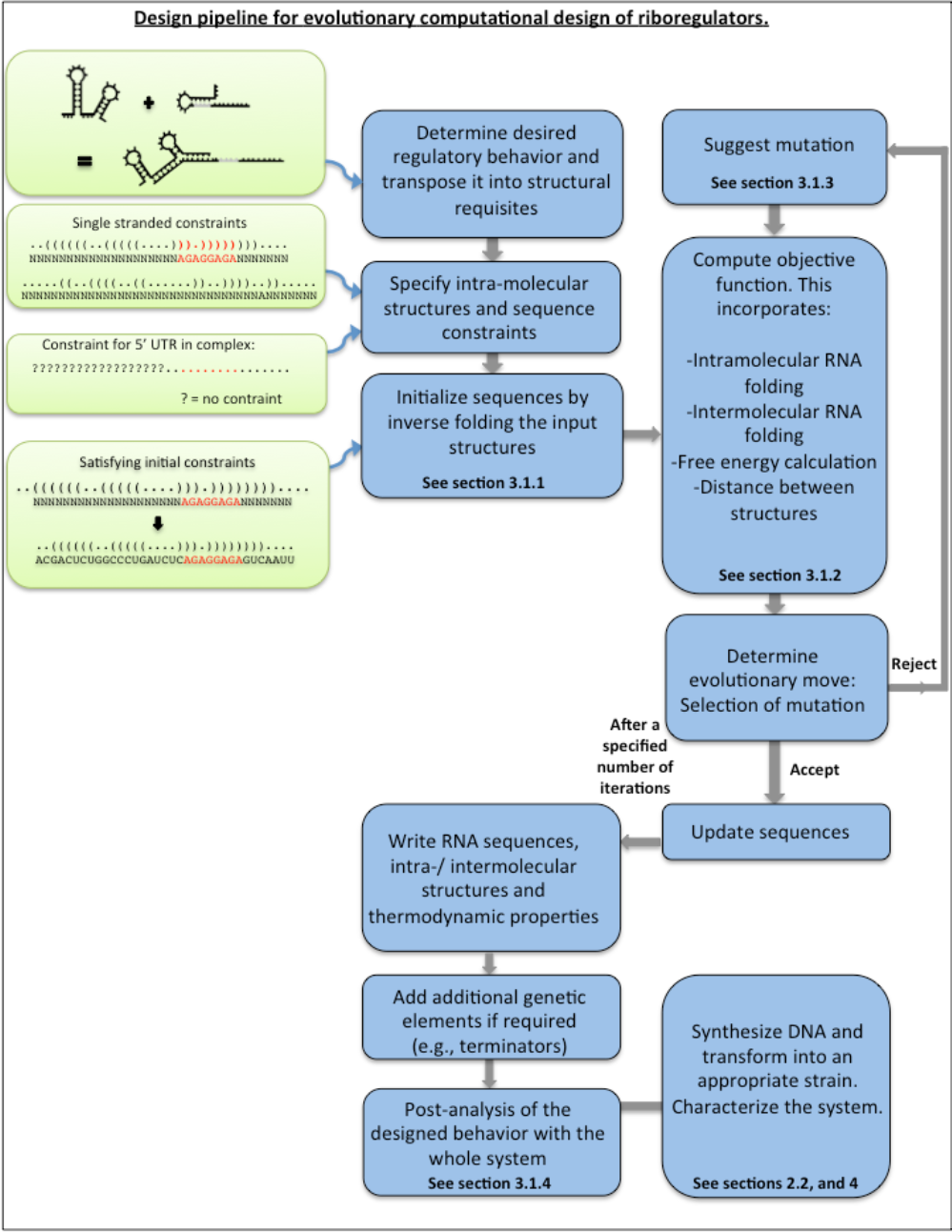
RNA circuit design pipeline. Automatic evolutionary design uses secondary structure scaffolds as a starting point to generate random starting RNAs. These are then sequencially mutated and selected to evolve them towards a solution which satisfies a user defined behavior.

#### 2.1.1 ViennaRNA (version 2.0)

1. Download the ViennaRNA package from http://www.tbi.univie.ac.at/∼ronny/RNA/index.html.
2. Follow the supplied instructions for installation. There is no need for additional libraries.
3. Use the following functions within the ViennaRNA package: (1) *fold*, to predict minimum free energy secondary structures, (2) *cofold*, as fold but for two different species, and (3) *inverse_fold*, to get sequences with predefined structures.

#### 2.1.2 Nupack (version 3.0)

1. Download the Nupack package from http://www.nupack.org/downloads/source.
2. Follow the supplied instructions for installation. There is no need for additional libraries.
3. Use the following functions within the Nupack package: (1) *concentrations*, to predict the equilibrium concentration of each species (single or complex) in a dilute solution given initial concentrations of the single ones, and (2) *subopt*, to determine all possible structures of the thermodynamic ensemble within an energy gap, and to check for structural robustness.

### 2.2 Necessary elements for *in vivo* implementation and expression

#### 2.2.1 Promoters

1. Use well-characterized constitutive or inducible promoters and choose them accordingly to the genetic background (see Table 2 for a list of useful promoters; see also Note 1).
2. It is essential that the chosen promoters have a known and characterized transcription start site (+1 position) to avoid truncation or alteration of the RNA sequence. Pay special care when using a promoter truncated to the transcription start site because the RNA sequence may affect the transcription initiation.

**Table 2:**
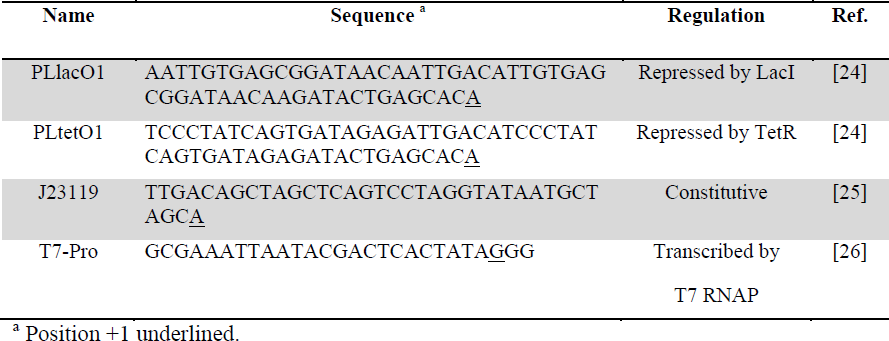
List of useful promoters for riboregulation.

#### 2.2.2 Transcription terminators

1. Transcription terminators are placed just downstream of the sRNA and of the coding sequence. Favor short rho-independent terminators with strong activity (i.e., with high free energy and long poly(U) tail; see Table 3 for a list of useful terminators in riboregulation).
2. Ideally, the computational design process should take into account the terminator sequence and structure, especially for the riboregulator. The terminator sequence can be specified as a sequence constraint. Alternatively, known toy models of transcription termination [29, 30] could be quantitative and predictive enough to be incorporated in the objective function.

**Table 3:**
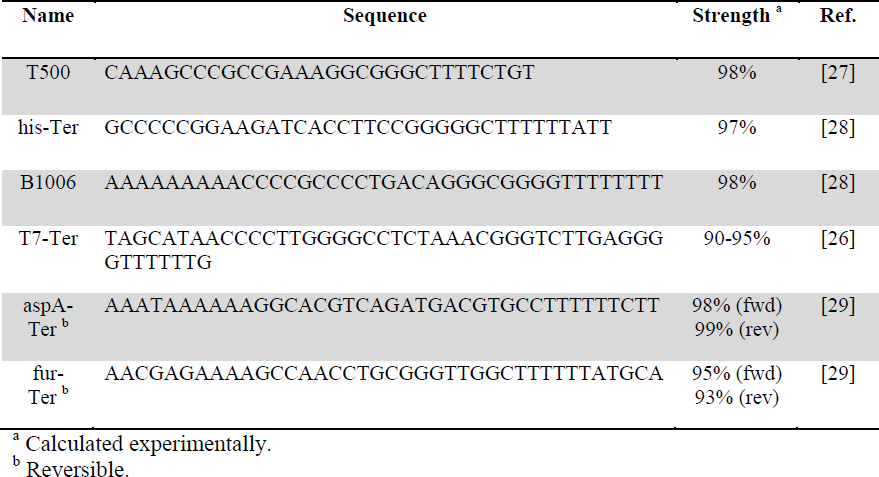
List of useful transcription terminators for riboregulation.

In addition, the expression of designed circuits in different genetic backgrounds depends on the characteristics of the engineered system and the functional properties one wants to study. To this end, several *E. coli* strains which are particularly suitable for RNA synthetic biology can be used (see Table 4).

**Table 4:**
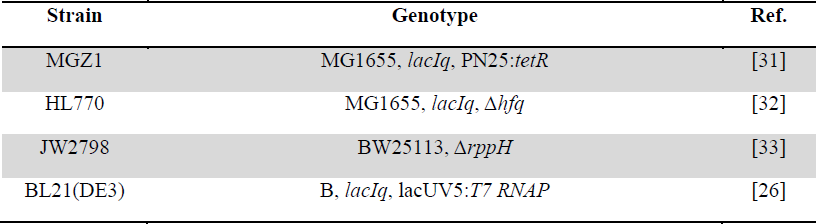
List of useful *E. coli* strains for studying riboregulation.

#### 2.2.3 Strains (cellular chassis)

1. The strain MGZ1 [30] is very useful for characterizing circuits. It constitutively expresses the transcription factors TetR and LacI (see also Note 2), enabling a control over promoters containing the relevant operators, like PLlacO1 and PLtetO1 [24]. Using IPTG and aTc, the level of expression of both RNA molecules can be finely tuned (see also Note 3).
2. In order to alter the function of an RNA system, various strains that are depleted in factors that are known to affect RNA *in vivo* can be used. These include: (1) RNase-deficient strains such as HT115 (Δ*rnc*, deleting RNase III) [34] or BL21 Star (mutated *rne*, making RNase E less efficient; Invitrogen),
3. (2) co-factor-deficient strains such as HL770 (Δ*hfq*) [32], or (3) strains deficient in RNA-processing proteins such as JW2798 (Δ*rppH*) [33].

#### 2.2.4 Plasmids

1. The RNA components can be together on one single plasmid, or separated on two different ones for co-transformation (see also Note 4).
2. Favor high copy number plasmids, with which the response is easier to detect due to concentration effects (presumably, the effective dissociation constant between two synthetic RNAs is high).

## 3 Methods (computational design)

In this section, we first describe the general strategy for implementing an automated design method. We formulate a combinatorial optimization problem to which we apply an evolutionary algorithm. In our case, this is based on Monte Carlo simulated annealing [35]. Then, we describe in detail how to use such an approach to design riboregulators acting, either as repressors or activators, on target 5’ UTRs (see Table 1 for experimentally verified examples). Finally, we describe ways to create more sophisticated systems.

### 3.1 General strategy of evolutionary design

Evolutionary design of synthetic RNA systems uses alternative rounds of computational assessment of RNA-RNA interaction followed by mutation [7]. Figure 1 outlines the general design pipeline. After initialization of the process, an objective function is used to evaluate the structures and free energies of RNAs, and a mutation operator modifies nucleotides or base pairs whilst keeping structural constraints for each species over the course of the evolutionary process. These rounds of mutation, scoring and selection are continued for a specified number of iterations or until a satisfactory solution is found. Afterwards, the sequences are output, and reviewed with an independent software, and assembled for characterization (see Fig. 2).

**Figure 2:**
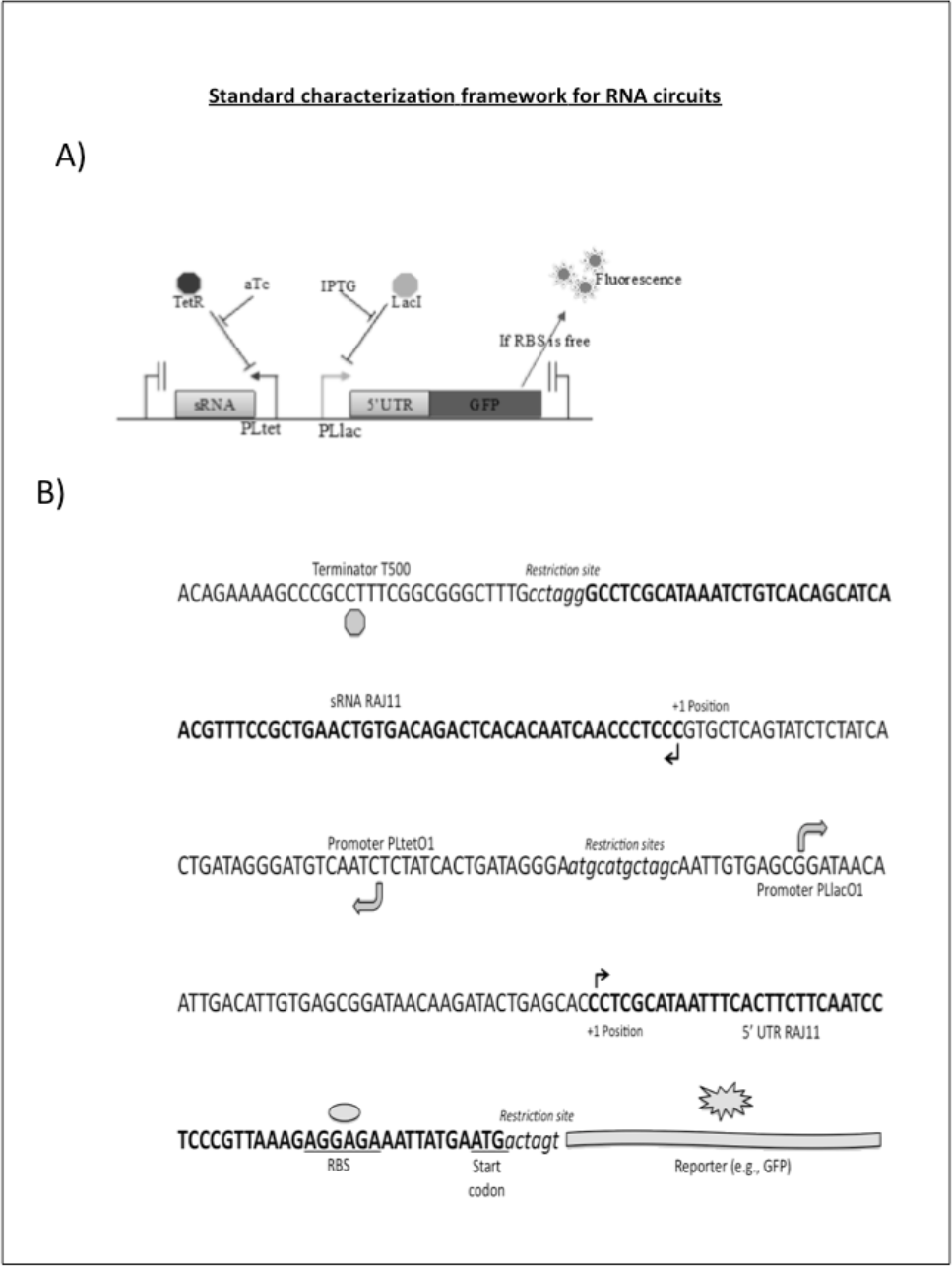
A) Characterization framework for an RNA circuit and B) Example of an assembled synthetic RNA device (here, the RAJ11 riboregulator and its 5’ UTR target controlling GFP), within an operon containing controllable promoters (PLlac and PLtet), terminators and with restriction sites for assembling controls or changing the reporter.

#### 3.1.1 Specifications and initiation

The desired regulatory behavior is converted into (intra-and intermolecular) structure specifications that can be read by a computer, coded in dot-bracket notation. For examples of specifications that can be used, see the sections 3.2 and 3.3 describing the design of negative and positive riboregulation, respectively. The definition of such structural specifications, as well as the sequence constraints, is a sensitive step, as the whole design process depends on it. The intramolecular structure specifications could be considered as constraints if desired, although this is not necessary, then the optimization of sequences would be performed over neutral networks in structure [7]. The free energies of hybridization and activation of the system and both the intra-and intermolecular structures of the 5’ UTR are used to evaluate our objective function.

1. Specify intramolecular structural specifications for each RNA species. The 5’ UTR and the sRNA can be fully constrained. Table 5 provides some usable structures, and natural or synthetic riboregulatory devices can inspire others.
2. Specify a conformational pattern for the complex species, where the ribosome-binding site (RBS) should be the only region subject to structural specifications, whilst the rest of the sequence of the 5’ UTR and the whole sRNA should have a large degree of freedom.
3. Specify sequence constraints, by defining nucleotides that are not allowed to mutate during the evolutionary process. This is essential in order to ensure that important RNA motifs such as RBSs or aptamers are not mutated.
4. Optionally, the sequence of one species can be fully constrained, and the system can be left to evolve the other RNA towards the intended regulation (constrained design).
5. Once all sequence and structure constraints are defined, use the inverse folding routine to find the initial, random sequences satisfying them.

**Table 5:**
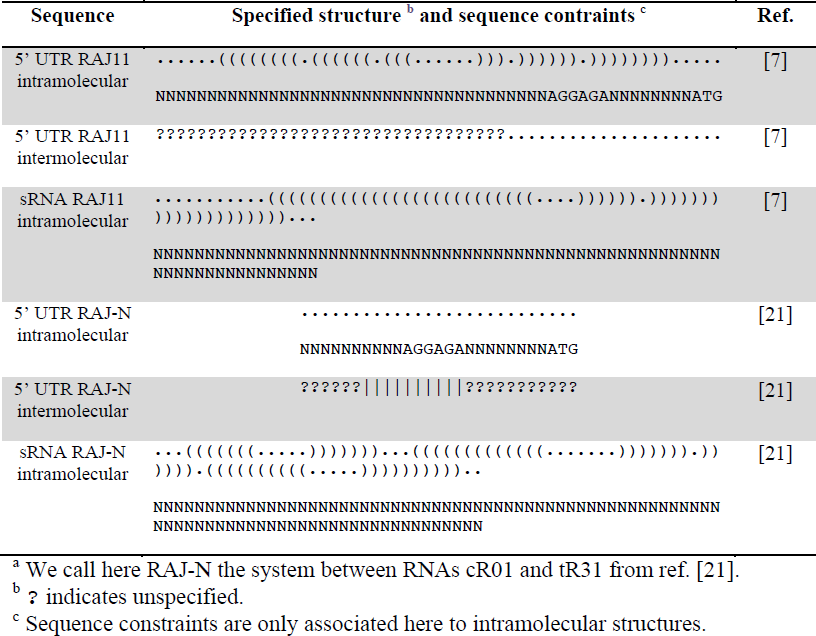
Examples of structure and sequence specifications for designing positive (system RAJ11) and negative (system RAJ-N ^a^) riboregulation.

#### 3.1.2 Selection

1. Build an objective function that can score a given RNA system taking into account the free energy of each species in the system as well as their secondary structures. Typically, this function is constructed with three weighted terms. The first term is the hybridization energy between the two RNAs. The second term is the activation energy, based on the exposed nucleotides of the seed region, which initiate the reaction [7]. The third is a structural term that accounts for the distance between the targeted structures and the actual ones (see sections 3.2 and 3.3 for examples).
2. The hybridization energy is the free energy release due to the RNA-RNA interaction. The energy gap should be as high as possible to ensure interaction *in vivo* (it is estimated at −17 Kcal/mol with Nupack for the system RAJ11 [7]).
3. The activation energy is given by the free energy release due to the interaction between the seed regions (six nucleotides for the system RAJ11 [7]). These regions must be unpaired in the single stranded forms so that the interaction can take place (see also Note 5).
4. The degree of exposition or blockage of the RBS within the secondary structure of the 5’ UTR (i.e., whether the Shine-Dalgarno box and surrounding nucleotides are paired or not) serves as the variable that accounts for function (such as protein translation).

#### 3.1.3 Mutation operator

1. The mutation operator takes both sequences as input, the riboregulator and 5’ UTR, and randomly mutates one of them. Nucleotides that are specified to be fixed (e.g., RBS) are not mutated. The mutation operator can make two types of mutation: Single-point mutations, or directed mutations.
2. For a single-point mutation, it chooses a nucleotide randomly. If this nucleotide is structurally unconstrained or unpaired at the intramolecular level, this operator simply mutates it to another one. If this nucleotide is constrained and paired intramolecularly, it mutates the base pair in order to keep secondary structure (see also Note 6).
3. Directed mutations are used to accelerate the search. The operator picks a set of consecutive nucleotides in one sequence (usually between 2 and 4 bases long), and introduces its reverse complement randomly in other sequence. As before, the operator assesses the structural constraints. From 50% to 90% of mutations can be directed.
4. The mutation operator refuses any mutation that creates a sequence of four or more identical nucleotides in a row.

#### 3.1.4 Post-analysis

In our settings, not all computational jobs converged to good solutions, and screening with a cutoff value of the objective function was required. In addition, false positives may be generated because the process is completely unsupervised. A post-analysis of good solutions (according to ViennaRNA and for the objective function defined) should therefore be performed, using the Nupack software and criteria that were not specified originally.

1. Analyze the base pair probabilities of key regions (typically the RBS) for the intra-and intermolecular folding states using the partition function. During the evolutionary design, the algorithm only considers the minimal free energy structure for simplicity and for reducing computation time, although in fact there is an ensemble of structures [17]. If the percentage of structures in the ensemble satisfying the structural specifications is lower than 90%, the system should be rejected.
2. Check the behavior of the system at equilibrium with Nupack for defined concentrations of 1 µM for each single species. If the complex species does not represent at least 95% of total at the equilibrium, the system should be rejected.
3. If it is not included in the design process, the terminator should be added to the final sequence of the sRNA before post-analysis (see also Note 7).
4. If the evolutionary process selects for structures that are only partially constrained, favor designs that have similar interaction patterns to those of known natural regulatory RNAs.

### 3.2 Negative riboregulation

Negative riboregulators have the ability to reduce protein expression of a target gene by either acting on its transcription rate (CRISPRi) [15], the mRNA stability (sRNA-induced degradation), or its translation rate. In this last case, the sRNA interacts with the 5’ UTR to block the RBS (and sometimes the start codon), by direct intermolecular interaction or by inducing a conformational change that results in intramolecular trapping, hiding it from the ribosome (see Fig. 3A) [21]. Table 5 shows structural specifications to design negative riboregulators.

1. The single stranded structure of the 5’ UTR must have an unpaired RBS.
2. Within the complex, the RBS must be paired (inter-or intramolecularly).
3. The predicted hybridization energy should be lower than −20 Kcal/mol.
4. Favor intramolecular trapping of the RBS to avoid unwanted cross-interaction.

**Figure 3:**
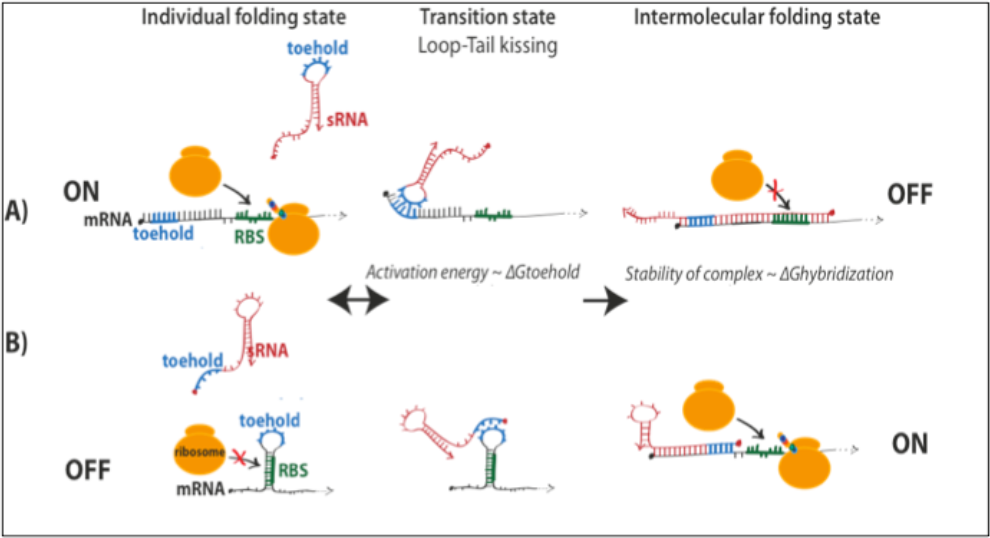
Mechanisms of: A) Negative riboregulators, where the default state is ON. The binding of sRNA blocks the RBS and represses translation; and of B) Positive riboregulators, where the default state is OFF, due to the RBS being hidden by the 5’UTR secondary of the mRNA. The binding of sRNA unfolds this structure, exposing the RBS and allowing translation. In both cases, the binding of the sRNA goes through a transition phase, when the toeholds — exposed regions of the RNAs — interact, followed by a full hybridization between the complementary regions. The length of the toeholds fixes the activation energy — ΔGtoehold – which determines the speed of the reaction, whereas the stability of the complex – ΔGhybridization – determines the equilibrium of the reaction.

### 3.3 Positive riboregulation

Positive riboregulators have the ability to increase protein expression of a target gene by acting on its transcription rate (with a fusion between the omega subunit of the RNA polymerase and a Cas9 nuclease) [16], or its translation rate. In this last case, the RBS in the target 5’ UTR (single stranded form) is *cis*-repressed, whilst the interaction with the sRNA causes a conformational change that releases the RBS and allows translation (see Fig. 3B). Table 5 shows structural specifications to design positive riboregulators.

1. The single stranded structure of the 5’ UTR must have a paired RBS.
2. Within the complex, the RBS must be unpaired.
3. The predicted hybridization energy should to be lower than −15 Kcal/mol.
4. Favor intermolecular interactions between different regions of the sRNA molecule.

### 3.4 A generic methodology for the design of logic RNA devices

In addition to its use for designing positive and negative riboregulators, the described methodology can be applied to optimize further structure-based regulatory specifications. This enables the design of RNA systems based on a larger diversity of mechanisms. Some strategies are described here.

#### 3.4.1 Three-input logic systems

The *de novo* design of RNA systems based on more than two RNA species can be challenging due to the exponential increase of the size of the search space. However, by either running the program on high-performance computers or by constraining the sequence space, convergence can be achieved. Indeed, in our recent work [21], we presented different examples of interaction mechanisms between two sRNAs and one 5’ UTR.

#### 3.4.2 Pseudo-3D modeling

One current limitation of automated RNA design can be the number of false positives produced, that is, sequences that behave as desired computationally but that do not work at the experimental level. Certainly, this is due to the lack of a comprehensive intermolecular interaction model (see also Note 8). It is expected that the incorporation of more structural and functional parameters into the objective function would increase the proportion of good designs. Although full 3D modeling would require too much computing task to be efficiently implemented in an evolutionary algorithm, a library of RNA 3D motifs and non-canonical base pairing, not restricted to the Watson-Crick model, could be used [36, 37].

#### 3.4.3 Integration of aptamer and ribozyme sequences

The possibility of creating modular and signal-processing RNA circuits with various kinds of inputs is of great interest. Aptamers are powerful examples of sensing modules that can be combined with actuator modules. They are hard to design *de novo* using energy models, as their function is mainly derived from their 3D structure. However, they can be exploited to design aptamer-based riboregulators, as shown in recent experiments with the theophylline aptamer [38, 39]. In addition, it is also possible to incorporate ribozymes or aptazymes as functional elements that can be rearranged in different RNA contexts [12]. This opens up the way for computer-assisted engineering of pathways that have RNA molecules as an input and as an output.

#### 3.4.4 Integration with CRISPR systems

In bacteria, engineered sRNAs based on the CRISPR-Cas system [14–16] have recently received a lot of attention, and have been harnessed for chromosome engineering [40] as well as regulation at the transcriptional [15, 16] and translational level [14]. Since an sRNA guides the Cas9 nuclease, this opens up the possibility for interaction with designed riboregulators. Indeed, we could design riboregulators to target the Cas9 recognition hairpin or the seed region of the guide RNA, leading to combinatorial RNA-mediated effects.

## 4 Notes

1. Recombination between homologous regions of promoters is a particularly frequent problem, and care must be taken when selecting promoters to avoid repetitions, symmetrical operators and other sequences prone to recombination [41].
2. Other strains that have the Z1 cassette integrated onto the genome are also available, such as DH5αZ1. However, in our experience, the MGZ1 cells are larger than DH5αZ1 cells, making them more suitable for microfluidics characterization. DH5αZ1 cells are also known to have a relatively low growth rate, which is why we suggest favoring the MGZ1 strain.
3. Bacterial growth rate must be taken into account for the analysis of the characterization data of the riboregulatory systems [42]. Translation rate, and not only protein expression, should be reported. This will avoid eventual artifacts produced by toxic inducers (such as aTc or theophylline), as protein expression is inversely proportional to growth rate, unless it has a degradation tag.
4. Riboregulators tend to work slightly better when both components are present on the same plasmid rather then when they are spread out on two different ones. This is presumably due to intracellular RNA diffusion and unexpected degradation.
5. The seed region is critical for a proper RNA-RNA interaction, as it is responsible for the initiation of the interaction. Mutations in this region can completely disrupt the regulatory behavior. On the other hand, this fact can be exploited to design orthogonal systems easily [6, 8].
6. The enforcement of a given structure for the two single species constrains the sequence space of possible solutions [7, 21]. By leaving those structures unconstrained, we could perform additions and/or deletions (not only replacements) of nucleotides during the optimization, and could potentially speed up the search of a solution.
7. In the case of the mRNA, the terminator and 5’ UTR are separated and isolated by the coding sequence and are unlikely to interact. For the sRNA, however, the terminator often represents a very significant proportion of the molecule (e.g., with the sequences provided in Table 3, the terminator would represent 20 to 40% of the final RNA). It is therefore essential to account for this, using preferentially short and stable ones.
8. Computational models do not account for RNA chaperones (e.g., Hfq) [43], nor for co-factors such as Mg^2+^ or Zn^2+^, which might have an impact on the designs. Moreover, the kinetics of RNA folding, binding, and turnover will have significant impact on the performance of designed RNA circuits [6, 12], and not only the thermodynamic properties.

## Acknowledgements

Work supported by the FP7-ICT-043338 (BACTOCOM) grant (to A.J.). We thank Anna Młynarczyk for critical reading of the manuscript. W.R. is supported by a DGA graduate fellowship, T.E.L by an AXA research fund PhD fellowship, and G.R. by an EMBO long-term fellowship co-funded by Marie Curie actions (ALTF-1177-2011).

## References

1. Waters, L.S., & Storz, G. (2009) Regulatory RNAs in bacteria. Cell 136, 615–628.

2. Isaacs, F.J., Dwyer, D.J., & Collins, J.J. (2006) RNA synthetic biology. Nat. Biotechnol. 24, 545–554.

3. Liang, J.C., Bloom, R.J., & Smolke, C.D. (2011) Engineering biological systems with synthetic RNA molecules. Mol. Cell. 43, 915–926.

4. Isaacs, F.J., Dwyer, D.J., Ding, C., Pervouchine, D.D., Cantor, C.R., & Collins, J.J. (2004) Engineered riboregulators enable post-transcriptional control of gene expression. Nat. Biotechnol. 22, 841–847.

5. Bayer, T.S., & Smolke, C.D. (2005) Programmable ligand-controlled riboregulators of eukaryotic gene expression. Nat. Biotechnol. 23, 337–343.

6. Lucks, J.B., Qi, L., Mutalik, V.K., Wang, D., & Arkin, A.P. (2011) Versatile RNA-sensing transcriptional regulators for engineering genetic networks. Proc. Natl. Acad. Sci. USA 108, 8617–8622.

7. Rodrigo, G., Landrain, T.E., & Jaramillo, A. (2012) De novo automated design of small RNA circuits for engineering synthetic riboregulation in living cells. Proc. Natl. Acad. Sci. USA 109, 15271–15276.

8. Mutalik, V.K., Qi, L., Guimaraes, J.C., Lucks, J.B., & Arkin, A.P. (2012) Rationally designed families of orthogonal RNA regulators of translation. Nat. Chem. Biol. 8, 447–454.

9. Na, D., Yoo, S.M., Chung, H., Park, H., Park, J.H., & Lee, S.Y. (2013) Metabolic engineering of Escherichia coli using synthetic small regulatory RNAs. Nat. Biotechnol. 31, 170–174.

10. Win, M.N., & Smolke, C.D. (2007) A modular and extensible RNA-based gene-regulatory platform for engineering cellular function. Proc. Natl. Acad. Sci. USA 104, 14283–14288.

11. Wieland, M., & Hartig, J.S. (2008) An improved aptazyme design and in vivo screening enable riboswitching in bacteria. Angew. Chem. Int. Ed. 47, 2604–2607.

12. Carothers, J.M., Goler, J.A., Juminaga, D., & Keasling, J.D. (2011) Model-driven engineering of RNA devices to quantitatively program gene expression. Science 334, 1716–1719.

13. Klauser, B., & Hartig, J.S. (2013) An engineered small RNA-mediated genetic switch based on a ribozyme expression platform. Nucleic Acids Res. 41, 5542–5552.

14. Qi, L., Haurwitz, R.E., Shao, W., Doudna, J.A., & Arkin, A.P. (2012) RNA processing enables predictable programming of gene expression. Nat. Biotechnol. 30, 1002–1006.

15. Qi, L.S., Larson, M.H., Gilbert, L.A., Doudna, J.A., Weissman, J.S., Arkin, A.P., & Lim, W.A. (2013) Repurposing CRISPR as an RNA-guided platform for sequence-specific control of gene expression. Cell 152, 1173–1183.

16. Bikard, D., Jiang, W., Samai, P., Hochschild, A., Zhang, F., & Marraffini, L.A. (2013) Programmable repression and activation of bacterial gene expression using an engineered CRISPR-Cas system. Nucleic Acids Res. doi: 10.1093/nar/gkt520.

17. McCaskill, J.M. (1990) The equilibrium partition function and base pair binding probabilities for RNA secondary structure. Biopolymers 29, 1109–1119.

18. Mathews, D.H., Sabina, J., Zuker, M., & Turner, D.H. (1999) Expanded sequence dependence of thermodynamic parameters improves prediction of RNA secondary structure. J. Mol. Biol. 288, 911–940.

19. Rodrigo, G., Carrera, J., Landrain, T.E., & Jaramillo, A. (2012) Perspectives on the automatic design of regulatory systems for synthetic biology. FEBS Lett. 586, 2037–2042.

20. Foster, J.A. (2001) Evolutionary computation. Nat. Rev. Genet. 2, 428–436.

21. Rodrigo, G., Landrain, T.E., Majer, E., Daròs, J.A., & Jaramillo, A. (2013) Full design automation of multi-state RNA devices to program gene expression using energy-based optimization. PLoS Comput. Biol. 9, e1003172.

22. Hofacker, I.L., Fontana, W., Stadler, P.F., Bonhoeffer, L.S., Tacker, M., & Schuster, P. (1994) Fast folding and comparison of RNA secondary structures. Monatsch. Chem. 125, 167–188.

23. Dirks, R.M., Bois, J.S., Schaeffer, J.M., Winfree, E., & Pierce, N.A. (2007) Thermodynamic analysis of interacting nucleic acid strands. SIAM Rev. 49, 65–88.

24. Lutz, R., & Bujard, H. (1997) Independent and tight regulation of transcriptional units in Escherichia coli via the LacR/O, the TetR/O and AraC/I1-I2 regulatory elements. Nucleic Acids Res. 25, 1203–1210.

25. Registry of Standard Biological Parts, MIT. http://parts.igem.org.

26. Studier, F.W., Rosenberg, A.H., Dunn, J.J., & Dubendorff, J.W. (1990) Use of T7 RNA polymerase to direct expression of cloned genes. Methods Enzymol. 185, 60–89.

27. Larson, M.H., Greenleaf, W.J., Landick, R., & Block, S.M. (2008) Applied force reveals mechanistic and energetic details of transcription termination. Cell 132, 971–982.

28. Cambray, G., Guimaraes, J.C., Mutalik, V.K., Lam, C., Mai, Q.A., Thimmaiah, T., Carothers, J.M., Arkin, A.P., & Endy, D. (2013) Measurement and modeling of intrinsic transcription terminators. Nucleic Acids Res. 41, 5139–5148.

29. Chen, Y.J., Liu, P., Nielsen, A.A., Brophy, J.A., Clancy, K., Peterson, T., & Voigt, C.A. (2013) Characterization of 582 natural and synthetic terminators and quantification of their design constraints. Nat. Methods 10, 659–664.

30. D’Aubenton Carafa, Y., Brody, E., & Thermes, C., (1990) Prediction of rho-independent Escherichia coli transcription terminators. A statistical analysis of their RNA stem-loop structures. J. Mol. Biol. 216, 835–858.

31. Dunlop, M.J., Cox, R.S., 3rd, Levine, J.H., Murray, R.M., & Elowitz, M.B. (2008) Regulatory activity revealed by dynamic correlations in gene expression noise. Nat. Genet. 40, 1493–1498.

32. Hussein, R., & Lim, H.N. (2011) Disruption of small RNA signaling caused by competition for Hfq. Proc. Natl. Acad. Sci. USA 108, 1110–1115.

33. Baba, T., Ara, T., Hasegawa, M., Takai, Y., Okumura, Y., Baba, M., Datsenko, K.A., Tomita, M., Wanner, B.L., & Mori, H. (2006) Construction of Escherichia coli K-12 in-frame, single-gene knockout mutants: the Keio collection. Mol. Syst. Biol. 2, 2006.0008.

34. Takiff, H.E., Chen, S.M., & Court, D.L. (1989) Genetic analysis of the rnc operon of Escherichia coli. J. Bacteriol. 171, 2581–2590.

35. Kirkpatrick, S., Gelatt, C.D., & Vecchi, M.P. (1983) Optimization by simulated annealing. Science 220, 671–680.

36. Leontis, N.B., Stombaugh, J., & Westhof, E. (2002) The non-Watson-Crick base pairs and their associated isostericity matrices. Nucleic Acids Res. 30, 3497–3531.

37. Das, R., Karanicolas, J., & Baker, D. (2010) Atomic accuracy in predicting and designing noncanonical RNA structure. Nat. Methods 7, 291–294.

38. Bayer, T.S., & Smolke, C.D. (2005) Programmable, ligand-controlled riboregulators of eukaryotic gene expression. Nat. Biotechnol. 23, 337–343.

39. Qi, L., Lucks, J.B., Liu, C.C., Mutalik, V.K., & Arkin, A.P. (2012) Engineering naturally occurring trans-acting non-coding RNAs to sense molecular signals. Nucleic Acids Res. 40, 5775–5786.

40. Jiang, W., Bikard, D., Cox, D., Zhang, F., & Marraffini, L.A. (2013) RNA-guided editing of bacterial genomes using CRISPR-Cas systems. Nat. Biotechnol. 31, 233–239.

41. Romero, D., Martínez-Salazar, J., Ortiz, E., Rodríguez, C., & Valencia-Morales, E. (1999) Repeated sequences in bacterial chromosomes and plasmids: a glimpse from sequenced genomes. Res. Microbiol. 150, 735–743.

42. Klumpp, S., Zhang, Z., & Hwa, T. (2009) Growth rate-dependent global effects on gene expression in bacteria. Cell 139, 1366–1375.

43. Vogel, J., & Luisi, B.F. (2011) Hfq and its constellation of RNA. Nat. Rev. Microbiol. 9, 578–589.

